# BPIFB3 interacts with ARFGAP1 and TMED9 to regulate non-canonical autophagy and RNA virus infection

**DOI:** 10.1101/2020.07.16.207035

**Authors:** Azia S. Evans, Nicholas J. Lennemann, Carolyn B. Coyne

## Abstract

Autophagy is a degradative cellular pathway that targets cytoplasmic contents and organelles for turnover by the lysosome. Various autophagy pathways play key roles in the clearance of viral infections, and many families of viruses have developed unique methods for avoiding degradation. Some positive stranded RNA viruses, such as enteroviruses and flaviviruses, usurp the autophagic pathway to promote their own replication. We previously identified the endoplasmic reticulum-localized protein BPIFB3 as an important regulator of non-canonical autophagy that uniquely impacts the replication of enteroviruses and flaviviruses. Here, we find that many components of the canonical autophagy machinery are not required for BPIFB3-regulated autophagy and identify the host factors that facilitate its role in the replication of enteroviruses and flaviviruses. Using proximity-dependent biotinylation (BioID) followed by mass spectrometry, we identify ARFGAP1 and TMED9 as two cellular components that interact with BPIFB3 to regulate autophagy and viral replication. Importantly, our data demonstrate that non-canonical autophagy in mammalian cells can be controlled outside of the traditional pathway regulators and define the role of two proteins in BPIFB3-mediated non-canonical autophagy.

**Summary Statement:** BPIFB3 is a regulator of a non-canonical cellular autophagy pathway that impacts the replication of enteroviruses and flaviviruses. Here we define ARFGAP1 and TMED9 as essential components of this pathway.

## Introduction

Autophagy is a catabolic cellular process that is responsible for the degradation of many cellular proteins, lipids, and excess organelles. The most widely characterized form of autophagy, termed macroautophagy, functions under the control of cellular nutrient sensors, such as mammalian target of rapamycin complex 1 (mTORC1) and AMP-activated protein kinase (AMPK) to degrade unnecessary cytoplasmic contents by targeting to the lysosome (Bento et al., 2016; Glick et al., 2010). Following activation from upstream signals, the initiation of autophagosome nucleation is controlled by the ULK1 complex which recruits the PI3K complex to commence formation of the isolation membrane (Shibutani and Yoshimori, 2014). The isolation membrane continues to expand, relying on specific receptors to target and sequester material within the nascent autophagosome prior to budding. The autophagosome then matures, fusing with other vesicles such as endosomes to form amphisomes, until it fuses with the lysosome to degrade the contents of the vesicle (Reggiori and Ungermann, 2017). Traditionally, macroautophagy has been referred to as canonical autophagy due to the extensive characterization of upstream signaling components and downstream factors that are responsible for the formation and trafficking of the autophagosome. However, a variety of other forms of autophagy exist, including microautophagy, chaperone mediated autophagy, and organelle specific autophagy that remain to be characterized to the same degree as canonical macroautophagy (Bento et al., 2016).

Despite the extensive characterization of signaling components involved in the regulation of macroautophagy, the origin of membranes responsible for the formation of nascent autophagosomes remains unclear. Each organelle within the secretory pathway has been implicated as a source of membranes for autophagosome biogenesis in mammalian cells, demonstrating a more complex relationship between membrane supply and autophagosome formation than what has been established in yeast. While it has been widely accepted that the endoplasmic reticulum (ER) plays an essential role in autophagosome formation, many studies have sought to narrow down the subdomain of the ER responsible for autophagosome nucleation. ER exit sites, ER-plasma membrane contact sites, and ER-mitochondria associated membranes have all been described as essential sites for autophagosome formation (Biazik et al., 2015; Carlsson and Simonsen, 2015; Ge et al., 2017; Graef et al., 2013; Omari et al., 2018). Additional studies have also suggested that sites of COPI vesicle formation within the Golgi may play an important role in early stages of autophagosome formation (Razi et al., 2009). Despite these efforts to discern the sites and membrane features of autophagosome biogenesis, there still remains much variability and uncertainty surrounding this process. While the site of autophagosome formation likely has a significant impact on the cargo to be degraded, many questions remain. For example, does the site of isolation membrane nucleation also impact the type of autophagy (canonical versus an alternate noncanonical pathway)? These questions become more complex when thinking about the context of autophagy during viral infections, as there can be direct implications for the location of autophagosome formation and the targeted clearance of viral particles or membrane associated viral replication organelles. We previously identified that the specific turnover of ER membranes by reticulophagy is a highly antiviral pathway for certain viruses (Lennemann and Coyne, 2017), however the impact of other specific autophagy pathways on viral replication is less clear.

Previously, we reported that an ER-localized protein, bactericidal/permeability-increasing protein (BPI) fold-containing family B, member 3 (BPIFB3), regulates a non-canonical form of autophagy to control RNA virus infection, including members of the enterovirus and flavivirus families (Delorme-Axford et al., 2014; Evans et al., 2020). BPIFB3 belongs to the BPIFB family of proteins named for their homology to BPI, a secreted antimicrobial protein. Despite the high degree of predicted structural homology, BPIFB3 and other ER-localized members of the family, BPIFB2 and BPIFB6, are not secreted and remain associated with the cytoplasmic face of the ER, despite lacking transmembrane domains (Delorme-Axford et al., 2014). Furthermore, our prior studies have determined that both BPIFB3 and BPIFB6 play important roles in vesicle trafficking and regulation of both enteroviruses and flaviviruses, including coxsackievirus B (CVB), dengue virus (DENV), and Zika virus (ZIKV) (Delorme-Axford et al., 2014; Evans et al., 2020; Morosky et al., 2016).

Our previous characterization of BPIFB3 revealed that its RNAi-mediated silencing resulted in a striking enhancement of autophagy, while it’s over expression restricted autophagy (Delorme-Axford et al., 2014). This regulation of autophagy has been demonstrated to have important implications in regulating the infection of two distinct families of viruses, enteroviruses and flaviviruses. Depletion of BPIFB3 specifically enhances the replication capacity of the enterovirus CVB, while restricting the replication of the two flaviviruses DENV and ZIKV (Delorme-Axford et al., 2014; Evans et al., 2020). These disparate impacts on replication highlight key replication differences between the two families of viruses and their sensitivity to turnover by BPIFB3-regulated autophagy. Recently, we demonstrated that depletion of BPIFB3 not only impacts autophagic flux, but also disrupts ER integrity, which can be reversed by inhibition of ER turnover via RETREG1-dependent reticulophagy (Evans et al., 2020), suggesting that the depletion of BPIFB3 broadly affects multiple aspects of autophagy. However, the specific components that interact with BPIFB3 to elicit these effects remain unclear. In this study, we defined the interactome of BPIFB3 and characterized the role of specific interactions on autophagy and RNA virus infection. Using BioID followed by mass spectrometry, we identified two binding partners of BPIFB3, ARFGAP1 and TMED9, which we found regulate BPIFB3-specific autophagy. Our data presented here point to a mechanism whereby BPIFB3 interaction with ARFGAP1 and TMED9 regulates the initiation of a form of non-canonical autophagy.

## Results

### BPIFB3 depletion-induced autophagy does not rely on canonical autophagy regulators

Given that BPIFB3 regulates a non-canonical autophagy pathway, we sought to characterize which components of the autophagic initiation machinery were required for the induction of autophagy during BPIFB3 depletion. To do this, we used two distinct viruses, CVB and DENV, as read outs for the reversal of autophagy induction given that our previous studies showed that BPIFB3-induced autophagy increases the replication of CVB, while it restricts DENV infection (Delorme-Axford et al., 2014; Evans et al., 2020). We tested components associated with three key phases of autophagy induction—autophagosome nucleation (ULK1), isolation membrane formation (the catalytic component of PI3K (PI3KC3), UVRAG1, and BECN1), and processing of LC3 (ATG7). To determine if these factors were essential for the induction of autophagy observed during BPIFB3 knockdown, we depleted each component by RNAi-mediated silencing in human brain microvascular endothelial cells (HBMEC) either alone or during the context of BPIFB3 depletion and infected with CVB or DENV. Consistent with previous findings, BPIFB3 depletion alone resulted in an enhancement of CVB replication (**Figure 1A**) and a restriction of DENV replication (**Figure 1B**). Silencing of the autophagosome nucleation kinase ULK1 independently had minimal effect on both CVB and DENV (**Figures 1A and 1B**). Consistent with these results, when ULK1 was co-depleted with BPIFB3 there was no change in the effects of BPIFB3 silencing on CVB or DENV replication (**Figure 1A and 1B**). These data directly indicate that the enhancement of autophagy observed during BPIFB3 depletion is not reversed when ULK1 expression is knocked down. We performed these same analyses for factors associated with isolation membrane formation (**Figure 1C and 1D**) and LC3 processing (**Figure 1E and 1F**), using both CVB and DENV infection levels as an indication of the inhibition of autophagy. We found that none of the canonical autophagy components tested were able to inhibit BPIFB3si-induced autophagy. Intriguingly, the co-depletion of BPIFB3 with BECN1 seemed to exaggerate the effect of BPIFB3 depletion alone, suggesting some degree in similarity between the effects of BECN1 and BPIFB3 depletion on CVB and DENV. However, BECN1 depletion was still insufficient to inhibit the enhancement of autophagy observed during BPIFB3 silencing. Knockdown efficiency of the siRNAs was confirmed by RT-qPCR (**Figure S1**). These data provide a clear indication that the regulation of autophagy during BPIFB3 depletion is not dependent on macroautophagy regulatory components.

**Figure 1:**
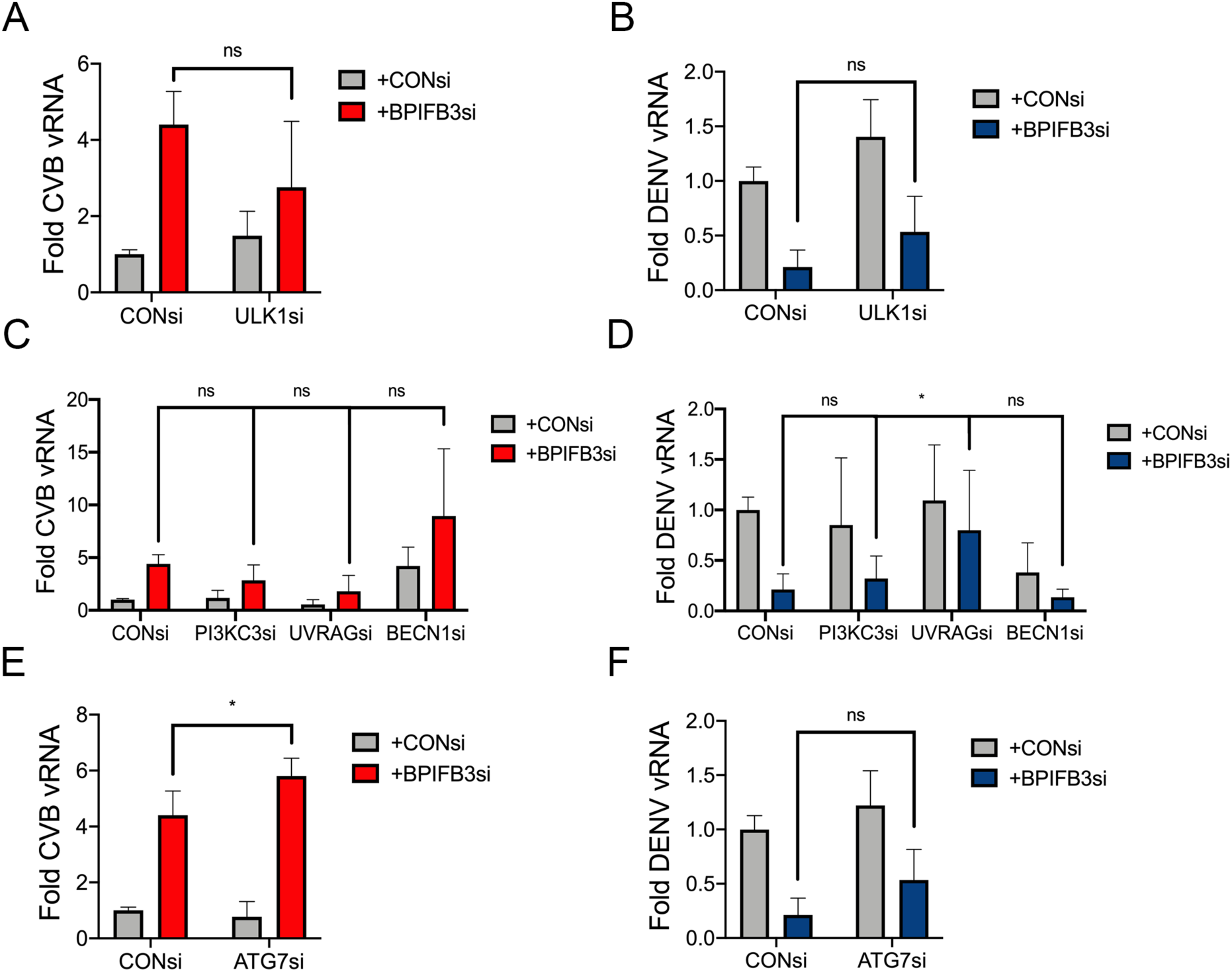
Canonical autophagy components are not required for BPIFB3si induced autophagy. HBMECs were depleted with BPIFB3 alone or in combination with key macroautophagy regulatory components, including factors involved in autophagosome nucleation, ULK1 **(A and B)**; components of the PI3K complex required for isolation membrane formation, PI3KC3, UVRAG, BECN1 **(C and D)**; and ATG7, which is required for LC3 and ATG5-ATG12 formation **(E and F)**. 48 hours post knockdown HBMEC were infected with CVB (16h) and DENV (48h) and viral replication was determined by RT-qPCR. CVB and DENV were used as a readout for BPIFB3si induced autophagy as our prior studies have determined BPIFB3 depletion increases CVB replication and restricts DENV infection. Data were analyzed using a two-way ANOVA, * P < 0.05.

### BioID mass spectrometry identifies ARFGAP1 and TMED9 as BPIFB3 interacting proteins

In order to identify proteins that interact with BPIFB3 and gain further insights into the mechanism(s) of its non-canonical autophagy regulation, we utilized the improved biotinylation based assay, BioID2 followed by mass spectrometry analysis (Kim et al., 2016). To achieve this, biotinylated proteins were isolated from HBMEC expressing BPIFB3-BioID2 or an empty BioID2 control vector. BPIFB3 interacting candidates were selected following mass spectrometry detection by analyzing raw peptide counts (**Figure 2A and 2D**) and percent protein coverage (**Figure 2B, 2C, and 2D**) as fold change to vector control samples. BioID identified a number of potential BPIFB3 interacting partners, including ARFGAP1 and TMED9 which have been previously implicated in the regulation of vesicle trafficking between the ER and Golgi (Beck et al., 2009; Pastor-Cantizano et al., 2016; Strating and Martens, 2009).

**Figure 2:**
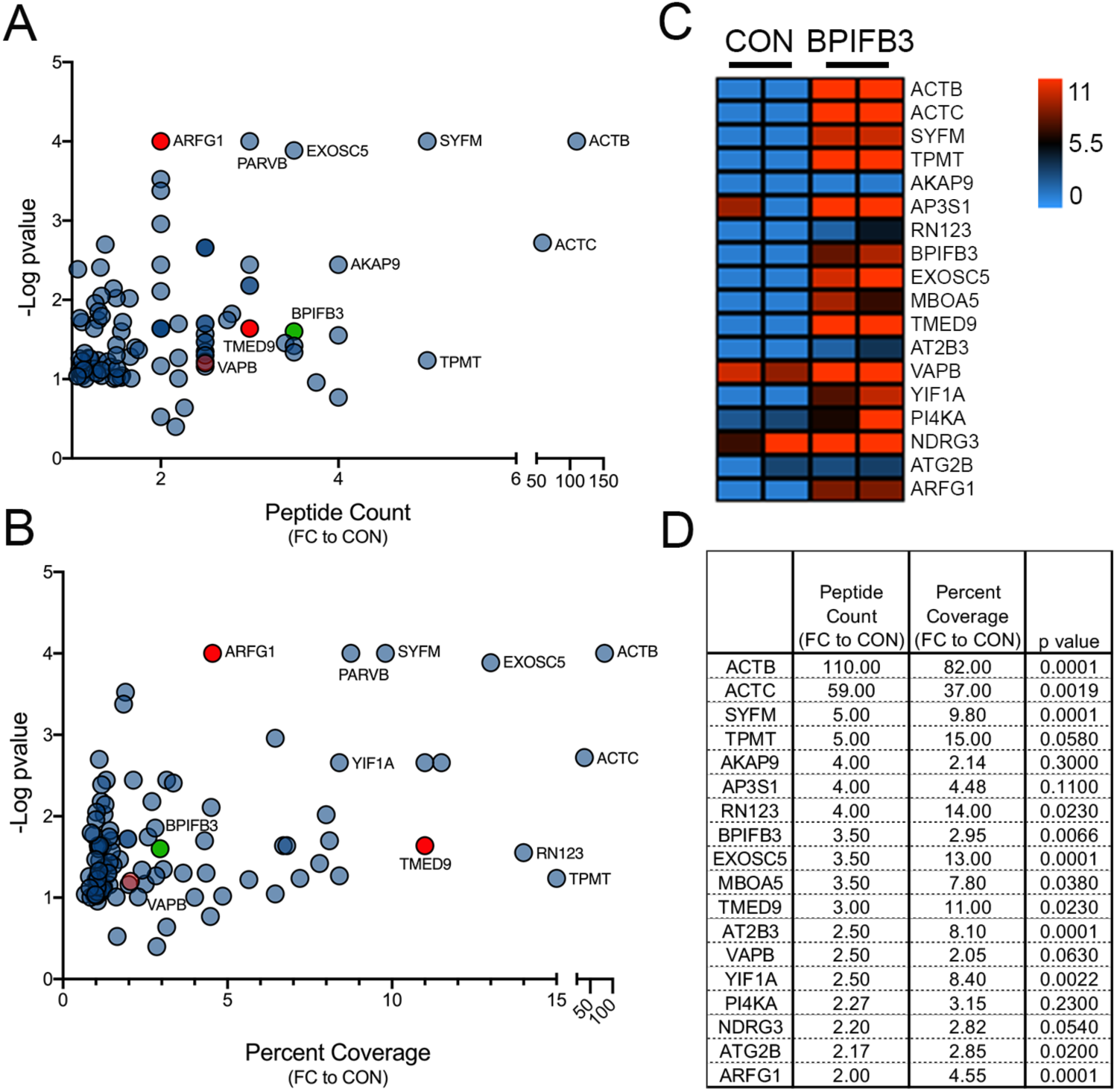
BioID analysis identified BPIFB3 interacting proteins. Mass spectrometry results from BioID tagged BPIFB3 expressed in HBMEC. Data represented as raw peptide counts (**A**) or percent protein coverage (**B**). (**C**) Heatmap showing individual percent coverage of mass spectrometry results for both vector control (CON) and BPIFB3. Average values for peptide counts, percent coverage, and significance for select candidates are shown in (**D**).

Validation of the BioID results by immunoprecipitation of BPIFB3 tagged with V5 expressed alone or with GFP-fused ARFGAP1 and TMED9 confirmed their interaction **(Figure 3A**). Interestingly, we also tested the interaction of BPIFB3 with VAPB, which was identified at low levels by the BioID screen (shown in **Figure 2A** and **2B** as the pale red point), and found that we were not able to validate its association with BPIFB3. These data confirm the specificity of interaction of BPIFB3 with ARFGAP1 and TMED9. We next examined the localization of ARFGAP1 and TMED9 when expressed with an ER marker or with BPIFB3. In agreement with our prior studies, ectopic expression of BPIFB3 with the ER marker KDEL-mCherry (**Figure 3B** top row) confirmed BPIFB3 localizes to the ER in a punctate expression pattern with a high Pearson’s correlation coefficient of 0.72 (**Figure 3D**). ARFGAP1 expression was primarily diffuse in the cytoplasm with a small amount of punctate perinuclear expression that is indicative of it’s known localization with the Golgi complex (Parnis et al., 2006). The punctate expression of ARFGAP1 further co-localized with the KDEL-mCherry marker (Pearson’s *r* = 0.82), as it has been previously reported that ARFGAP1 enhances both anterograde and retrograde trafficking between the ER and Golgi (**Figure 3B** middle row) (Dong et al., 2010; Shiba et al., 2011). Interestingly, co-expression of ARFGAP1 with BPIFB3 disrupted the punctate Golgi localization of ARFGAP1 (**Figure 3C** top row) however still remained highly co-locaized with BPIFB3 (Pearson’s *r* = 0.72) (**Figure 3D**). Expression of ARFGAP1 also altered the morphology of BPIFB3 localization, causing BPIFB3 to localize in a more reticular pattern with clusters of aggregated signal (**Figure 3C** top row). Ectopic expression of TMED9 was highly localized to the ER in a reticular pattern (Pearson’s *r* = 0.93), which is retained upon co-expression with BPIFB3 (**Figure 3B and 3C** bottom rows). Likewise, expression of TMED9 does not alter the localization pattern of BPIFB3. These results show that BPIFB3 interacts and colocalizes with ARFGAP1 and TMED9.

**Figure 3:**
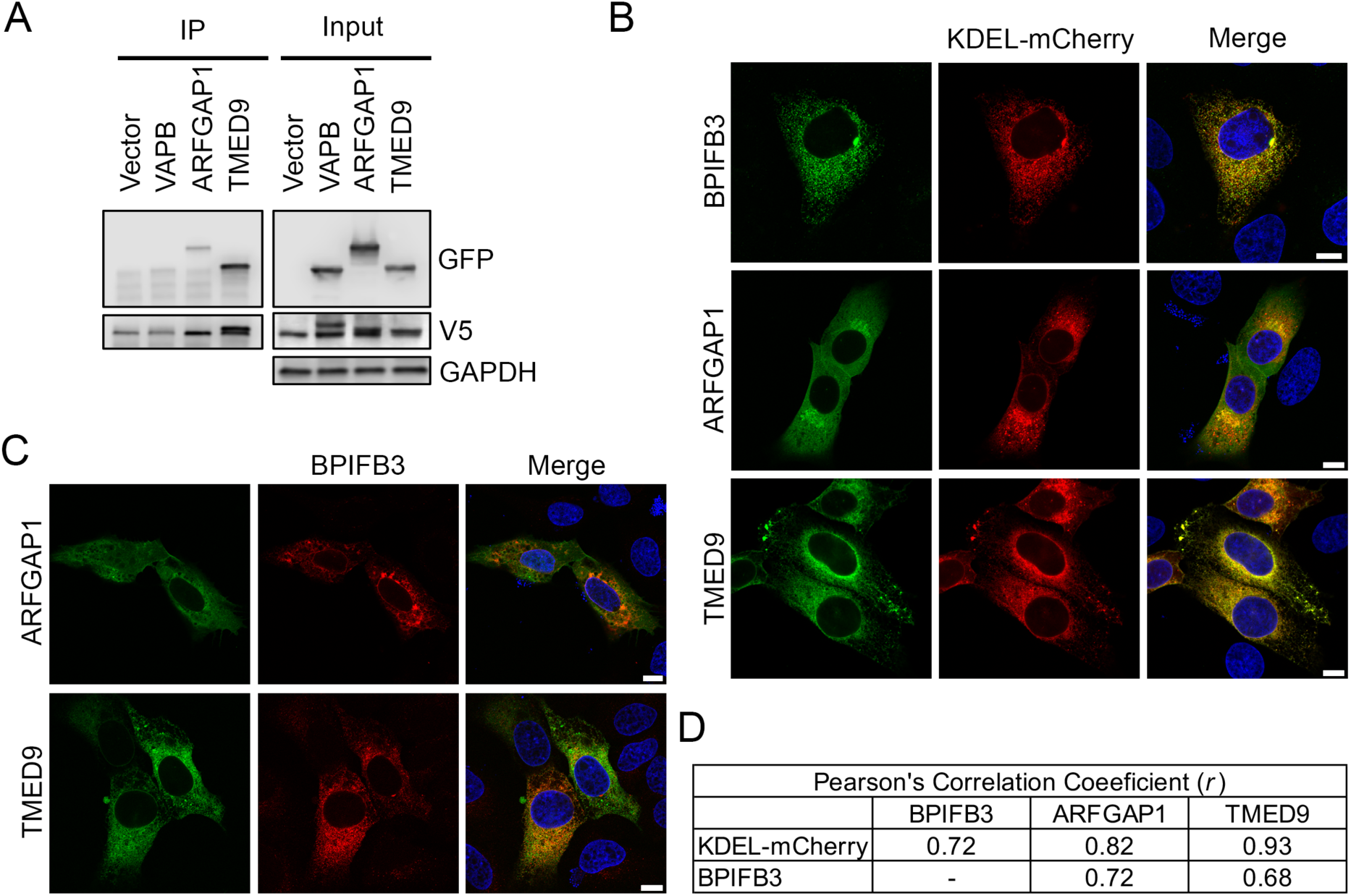
Validation of BioID results confirms ARFGAP1 and TMED9 interact with BPIFB3. (**A**) Immunoprecipitation of BPIFB3-V5 with select candidates identified by BioID, VAPB-GFP, ARFGAP1-GFP, and TMED9-GFP. Note that the double band on the V5 input blot is residual VAPB (top band) above BPIFB3 (bottom band) and is not a BPIFB3-V5 doublet. (**B**) Immunofluorescence imaging of U2OS cells expressing BPIFB3, ARFGAP1, and TMED9 (all green) with the ER marker, KDEL-mCherry. Scale bars are 10 μm. (**C**) Co-localization of BPIFB3 (red) was assessed with both ARFGAP1 and TMED9 (green) in U2OS cells by immunofluorescence. Scale bars are 10 μm. (**D**) A Pearson’s correlation analysis was used to determine colocalization between ARFGAP1, TMED9, and BPIFB3.

### ARFGAP1 and TMED9 expression are required for BPIFB3-mediated autophagy

Given that a hallmark of BPIFB3-silencing is increased levels of autophagy, we next sought to determine whether ARFGAP1 or TMED9 were required for this phenotype. HBMEC were transfected with a control siRNA (CONsi) or with BPIFB3si alone or in combination with siRNAs against ARFGAP1 and TMED9 (ARFGAP1si or TMED9si). Samples were then infected with CVB, which enhances the levels of BPIFB3si-induced autophagy (Delorme-Axford et al., 2014). Consistent with our prior studies, BPIFB3 silencing induced an increase in lipidated LC3 (LC3 II), which can be distinguished from non-lipidated LC3-I by immunoblot (**Figure 4A**). LC3-II levels were further enhanced in BPIFB3si-transfected cells infected with CVB (**Figure 4A**). Depletion of ARFGAP1 and TMED9 independently had minimal effects on LC3 levels during CVB infection (**Figure 4A**). However, co-depletion of ARFGAP1 and TMED9 with BPIFB3 led to a decrease in LC3-II levels compared to BPIFB3si alone, suggesting that the depletion of each inhibited BPIFB3si induced autophagy enhnacement. In contrast co-depletion of BPIFB3 with VAPB, did not reverse BPIFB3si induced autophagy (**Figure 4A**).

**Figure 4:**
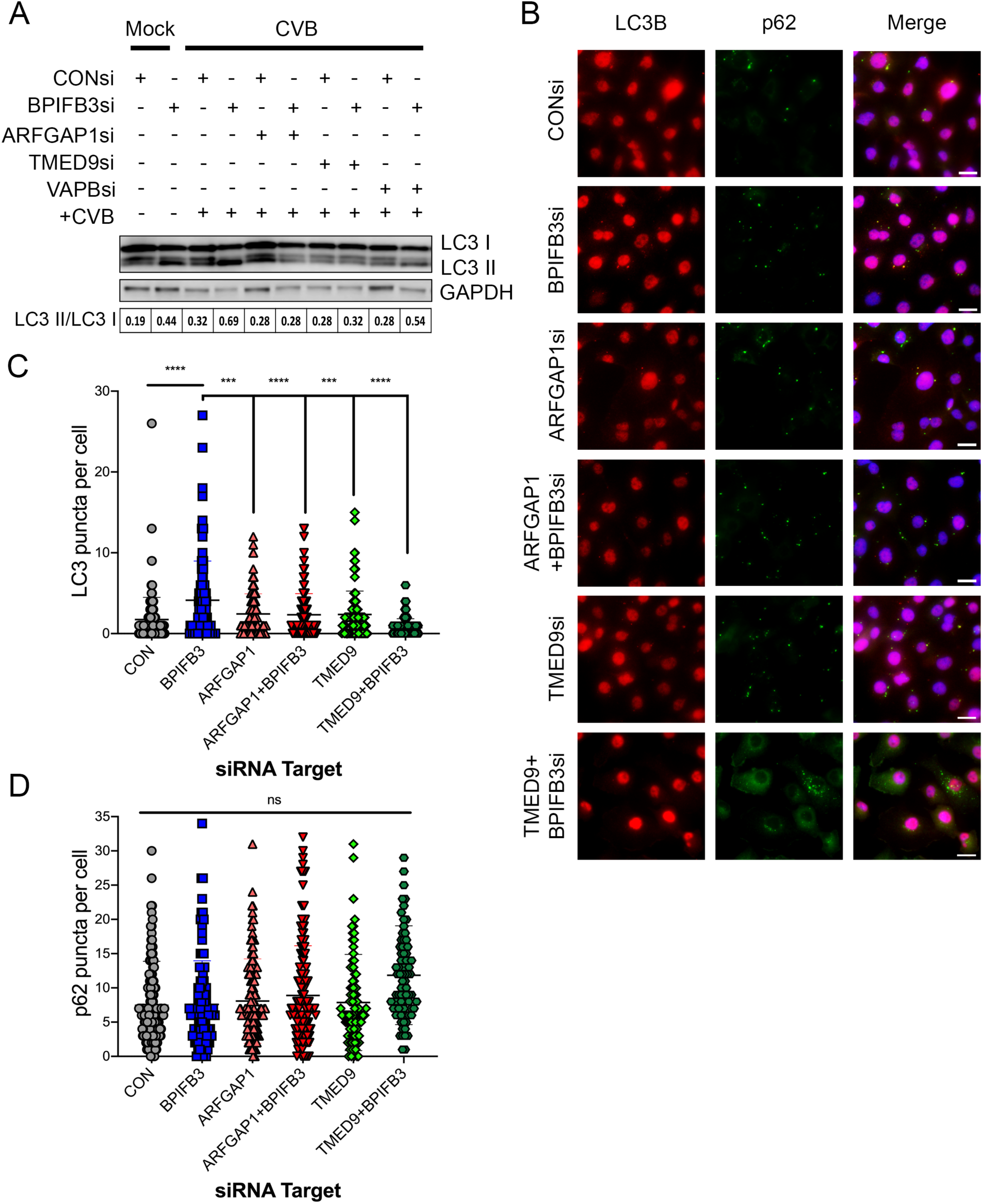
ARFGAP1 and TMED9 are required for BPIFB3si induced autophagy. (**A**) Western blot for LC3 protein levels from HBMEC transfected with BPIFB3si alone or in combination with ARFGAP1si, TMED9si, or VAPBsi. HBMEC were infected with CVB to exacerbate the effects of BPIFB3si induced autophagy and delineate changes in LC3 protein expression. The ratio of LC3 II/LC3 I shown in the table below, indicates the level of autophagy. (**B**) HBMECs depleted of BPIFB3 alone or in combination with ARFGAP1 or TMED9 were analyzed for level of autophagy by immunofluorescence imaging ot LC3 (red) and p62 (green). Scale bars are 25 μm. Quantification of LC3 and p62 puncta are shown in (**C and D**). Data were analyzed using a one-way ANOVA, *** P <0.001, **** P < 0.0001.

In order to confirm that ARFGAP1 and TMED9 depletion reversed the enhancement of autophagy in BPIFB3si-transfected cells, we used immunofluorescence-based microscopy to examine autophagy levels on a per cell basis. We performed these experiments in the absence of CVB infection to define the effect of BPIFB3 silencing independent of viral infection. LC3 positive autophagosomes and p62 positive vesicles were quantified in HBMEC transfected with CONsi or BPIFB3si alone or in combination with ARFGAP1 and TMED9 siRNAs **(Figure 4B-4D**). In agreement with our previous findings (Delorme-Axford et al., 2014; Evans et al., 2020), BPIFB3 depletion lead to an enhancement of LC3 positive puncta, which corresponds to the increase in lipidated LC3 observed by western blot. Silencing of ARFGAP1 and TMED9 alone had no impact on the number of LC3 positive autophagosomes when compared to control cells (CONsi). In contrast, co-depeltion of ARFGAP1 or TMED9 with BPIFB3 inhibited the induction of autophagy induced by BPIFB3si alone (**Figures 4B-4D**). Despite the clear changes observed in LC3 levels, it did not correspond with changes in the number of p62 positive puncta across any conditions, confirming our previosly published data that BPIFB3 regulates a non-canonical form of autophagy and not macroautophagy (Delorme-Axford et al., 2014; Evans et al., 2020).

To further define the impact of ARFGAP1 and TMED9 silencing on BPIFB3-associated autophagy, we performed transmission electron microscopy (TEM). Consistent with our previous work, BPIFB3 depletion increased the number of autophagy-associated vesicles (e.g. double membrane vesicles, amphisomes, and lysosomes) compared to control cells, which was reversed by co-depletion of either ARFGAP1 or TMED9 (**Figure 5A and 5B**). Depletion of ARFGAP1 alone lead to an increase in the number amphisomes present, indicative of the role that ARFGAP1 plays in both COPI and endocytic vesicle trafficking (Bai et al., 2011; Beck et al., 2009), however there was no observable impact on autophagy levels (**Figure 5B**). The co-depletion of ARFGAP1 with BPIFB3 showed a similar phenotype, with fewer autophagy associated vesicles than BPIFB3si alone (quantified in **Figure 5B**). Additionally, we also observed an increase in the presence of ER membranes throughout the cytoplasm during co-depletion of ARFGAP1 and BPIFB3 that was not apparent in the individual knockdowns. This phenotype shows interesting parallels to the effects of BPIFB3 silencing on ER morphology that we previously showed was rescued by the depletion of the reticulophagy receptor RETREG1 (Evans et al., 2020). The depletion of TMED9 alone showed no significant effect on the formation of autophagy-associated vesicles, but did show modest evidence of changes in ER morphology, with less continuous ER sheet-like membranes (**Figure 5A**). This is likely associated with the role TMED9 (also referred to p24α_2_) plays in ER exit site formation (Lavoie et al., 1999). In agreement with LC3 immunoblots and immunofluorescence, the co-depletion of TMED9 with BPIFB3 significantly reversed the increase in autophagy observed during BPIFB3 depletion alone (**Figure 5B**). Futhermore, we previously published that the depletion of BPIFB3 directly impacts ER morphology (Evans et al., 2020), and the TEM data presented here demonstrates not only that co-depletion of ARFGAP1 or TMED9 with BPIFB3 reverses the BPIFB3si-enhanced autophagy, but also further impacts ER morphology during BPIFB3 depletion.

**Figure 5:**
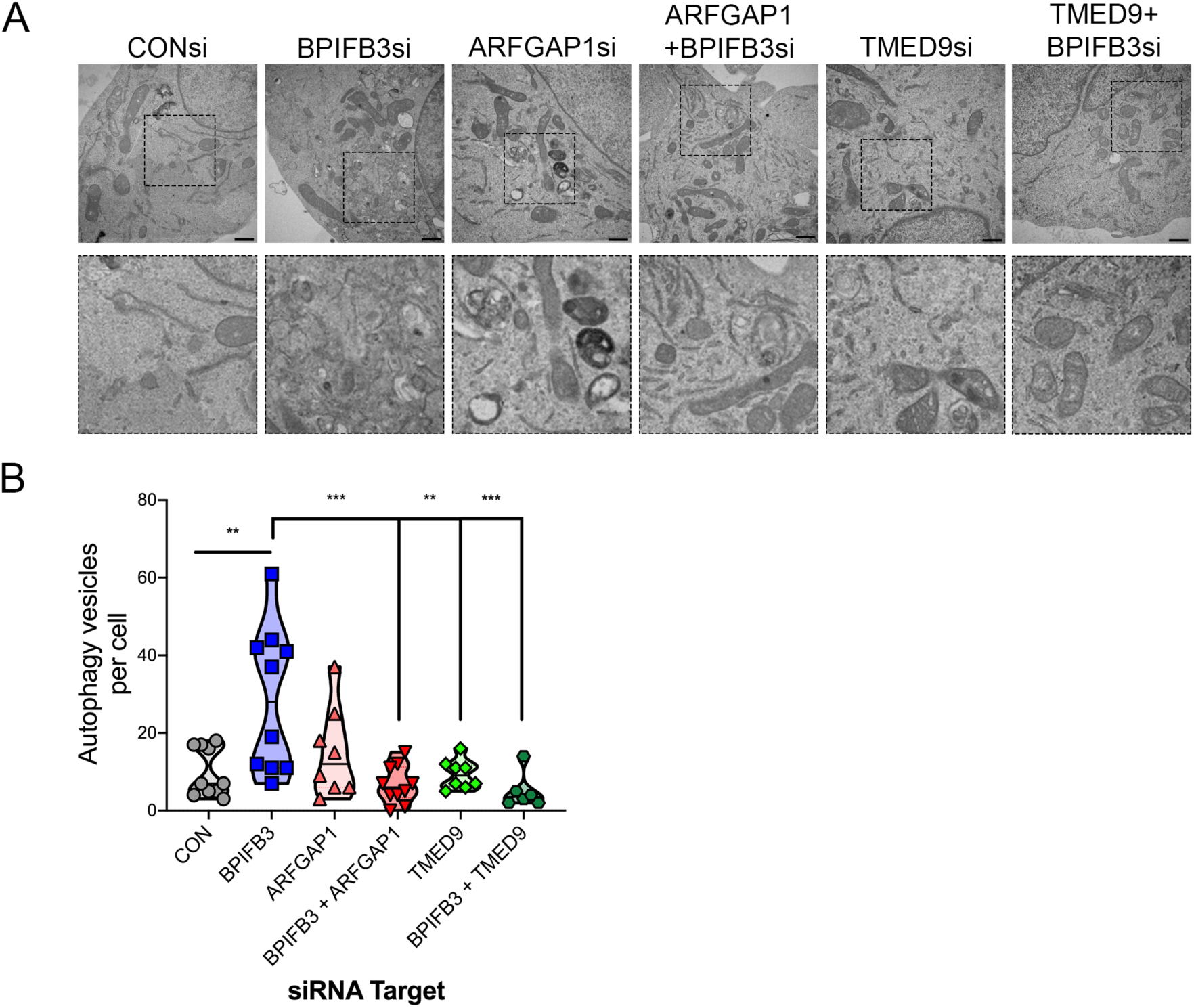
ARFGAP1 and TMED9 depletion reverse BPIFB3si morphology. (**A**) Transmission electron microscopy of HBMEC transfected with BPIFB3si alone or in combination with the BioID hits, ARFGAP1 and TMED9. Top row depicts large sections of the cytosol with dashed box around the zoomed in region shown in the bottom panel. Scale bars are 800 nm. (**B**) The number of autophagy-associated vesicles (e.g. double membrane vesicles, amphisomes, and lysosomes) were quantified per cell to determine if ARFGAP1 and TMED9 reversed the enhancement in non-canonical autophagy observed by BPIFB3 depletion. Data were analyzed using a one-way ANOVA, ** P <0.01, *** P < 0.001.

### Characterization of autophagy organelle markers during BPIFB3 depletion

Given that our characterization of key autophagy regulators on BPIFB3si induced autophagy showed no impact on the effects of autophagy induction (**Figure 1**), we performed immunofluorescence imaging of key organelle markers during the co-depletion of BPIFB3 with ARFGAP1 or TMED9. First, we examined lysosomal morphology by immunofluorescence microscopy of the late endosome/ lysosome marker LAMP1 (**Figure 6A**). In addition, samples were co-immunostained with the macroautophagy marker p62 to confirm that the increase in autophagy associated vesicles observed by TEM were not consistent with the macroautophagy pathway. In agreement with previous findings (Delorme-Axford et al., 2014; Evans et al., 2020) and our TEM data (**Figure 5**), BPIFB3 depletion increased the number of lysosomes, but not p62 positive vesicles (**Figure 6A**). Intriguingly, depletion of ARFGAP1 and TMED9 independently also increased the numbers of LAMP1 positive late endosomes/ lysosomes, which maycorrespond to the increased number of amphisomes observed by TEM. Despite this observable increase in LAMP1 vesicles during independent knockdown of each factor, the co-depletion of ARFGAP1 or TMED9 with BPIFB3 resulted in a reduction in the number of LAMP1 lysosomes compared to each of the single siRNA knockdowns (**Figure 6A**). These data confirm our TEM results that the co-depletion of ARFGAP1 and TMED9 with BPIFB3 reduces the number of autophagy associated vesicles beyond the autophagosome marker LC3 alone.

**Figure 6:**
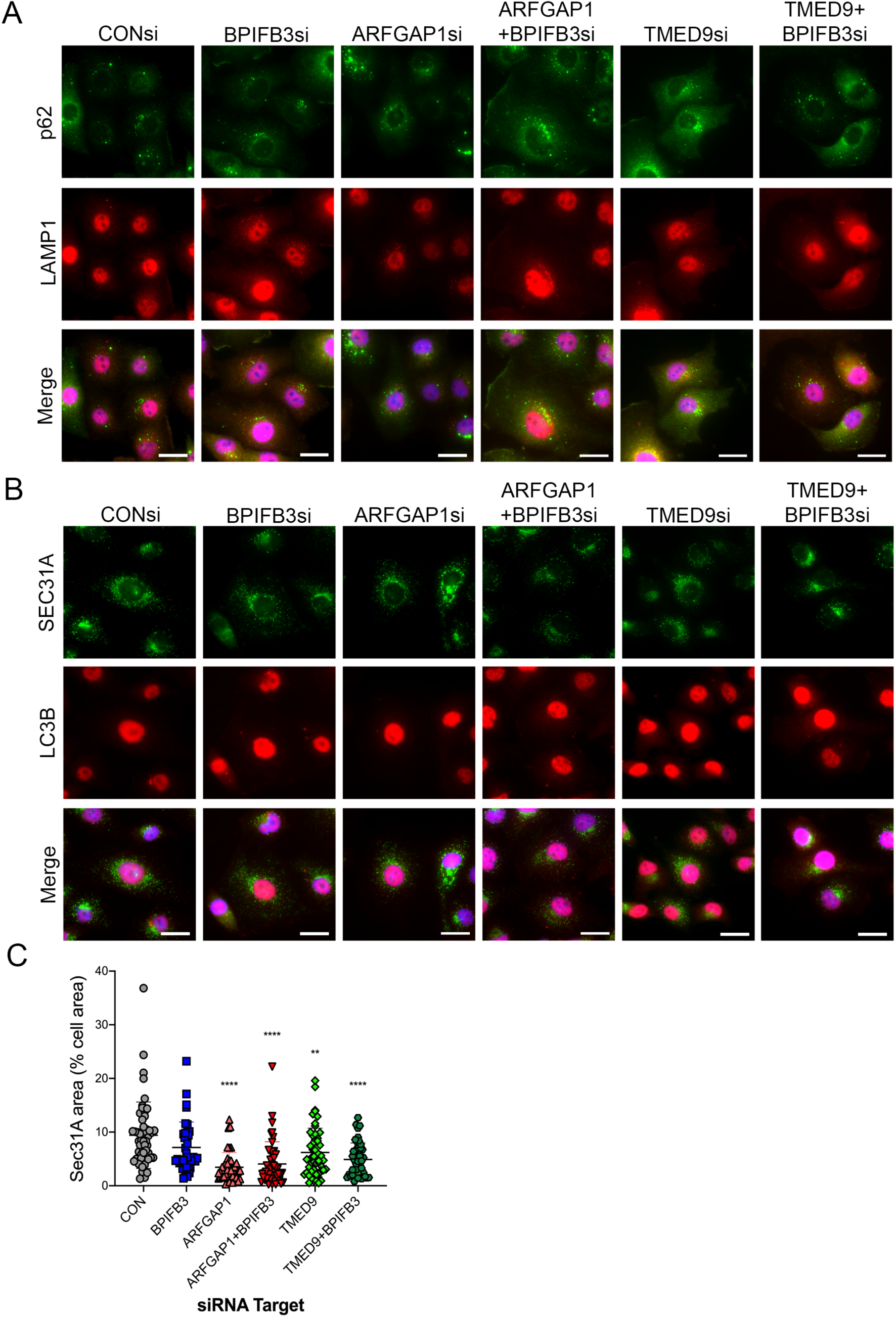
Characterization of autophagy related organelle markers during the co-depletion of BPIFB3 and ARFGAP1 or TMED9. (**A**) HBMEC transfected with each indicated combination of siRNAs were fixed and stained with antibodies to the macroautophagy receptor p62 and the late endosome/ lysosome marker LAMP1. (**B**) HBMEC transfected with BPIFB3si alone or in combination with ARFGAP1 or TMED9 were fixed and stained for the COPII vesicle coat component, Sec31A, and the autophagy marker, LC3B. Scale bars are 25 μm. (C) Quantification of Sec31A positive signal as percent of total cell area by siRNA knockdown conditions. Data were analyzed using a one-way ANOVA of each siRNA condition compared to control (CON), ** P <0.01, *** P < 0.001, **** P < 0.0001.

We also examined the morphology of ER exit sites during BPIFB3 depletion alone or with ARFGAP1 and TMED9. ARFGAP1 is a well-known regulator of COPI vesicle trafficking, while TMED9 has also been suggested to impact COPI trafficking (Beck et al., 2009; Lavoie et al., 1999; Pastor-Cantizano et al., 2016). However, our previous studies determined that BPIFB3 depletion has no effect on COPI morphology (Morosky et al., 2016). Despite its known role in COPI trafficking, early reports of TMED9 function suggest a role in ER exit site formation and morphology (Lavoie et al., 1999). Therefore we aimed to determine if the BPIFB3-ARFGAP1/TMED9 axis had a role in maintaining ER exit site morphology. We also sought to determine if ER exit sites served as sites of BPIFB3si-induced autophagosome formation. Immunofluorescence imaging of Sec31A, a component of the COPII coat complex, and LC3B during BPIFB3 depletion alone or with ARFGAP1 and TMED9 (**Figure 6B**), demonstrated no clear association between LC3 puncta and ER exit sites. Despite this, we observed morphological changes of Sec31A during ARFGAP1 and TMED9 depletion (**Figure 6B**). Quantification of Sec31A staining revealed a reduction in the total Sec31A area during ARFGAP1 and TMED9 depletion (**Figure 6C**). This reduction of ER exit site area was maintained during the co-depletion of ARFGAP1 and TMED9 with BPIFB3, but was not seen during BPIFB3 silencing alone. This suggests that ARFGAP1 and TMED9’s known role in COPI trafficking may extend to the regulation of ER exit site morphology. While there is no evidence of direct association between ER exit sites and BPIFB3si induced autophagy, these changes observed in ER exit site morphology might have downstream implications for the impact on vesicle trafficking and may explain the reversal autophagy that occurs.

To further characterize the role of ARFGAP1 and TMED9 during BPIFB3 knockdown, we ectopically expressed GFP-tagged ARFGAP1 or TMED9 in control or BPIFB3 depleted cells. Expression of ARFGAP1 during BPIFB3si conditions showed no changes of localization or morphology (**Figure 7A**). However, the expression of TMED9 during BPIFB3 depletion showed slight changes in distribution of its reticular pattern paired with increased co-localization with ER exit sites marked by Sec31A (**Figure 7B**). Taken together, these data indicate clear alterations in the COPI and COPII trafficking pathways during BPIFB3 depletion that is dependent on the presence of TMED9.

**Figure 7:**
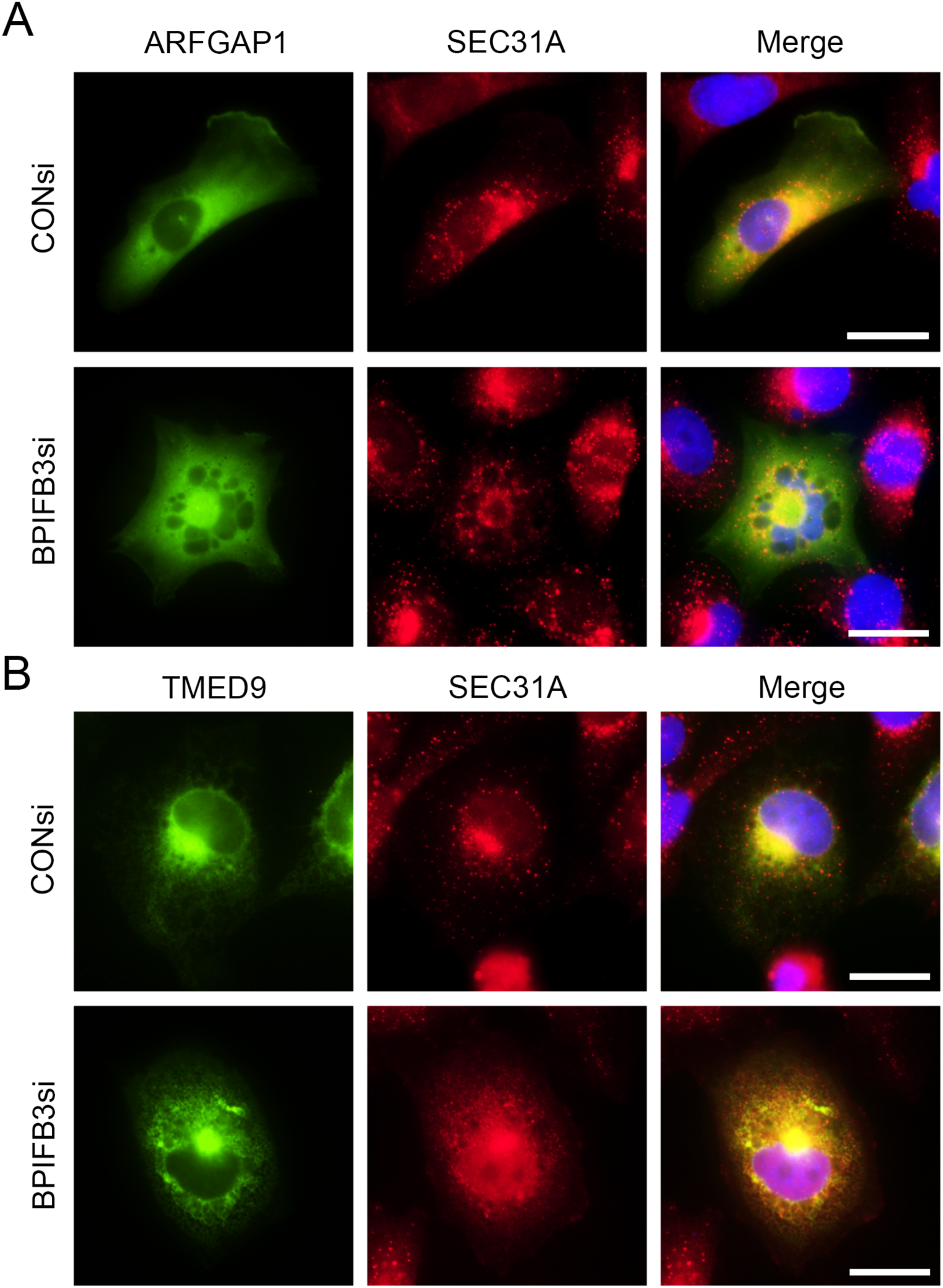
TMED9 expression effects ER exit site morphology during BPIFB3 depletion. CONsi or BPIFB3si HBMEC were transfected with ARFGAP1-GFP (**A**) or TMED9-GFP (**B**) and stained for the Sec31A. Scale bars are 25.

### ARFGAP1 and TMED9 are required for BPIFB3si modulation of RNA virus infection

Because we found that co-depletion of ARFGAP1 and TMED9 both reversed the effects of BPIFB3si-mediated autophagy, we next determined whether they were required for BPIFB3-mediated regulation of RNA virus infection Analysis of CVB (**Figure 8A**) and DENV (**Figure 8B**) infection upon depletion of ARFGAP1 or TMED9 had no effect on viral replication, whereas co-depletion with BPIFB3 demonstrated a reversal of the effects of BPIFB3 silencing alone. The knockdown efficiency of siRNAs targeting ARFGAP1 and TMED9 was confirmed by RT-qPCR (**Figure S2**). These data are in contrast to our findings with canonical regulators of autophagy (**Figure 1**). Given that ARFGAP1 and TMED9 reverse the phenotypes associated with each virus, confirms our findings that the effects of BPIFB3si are fully reversed.

**Figure 8:**
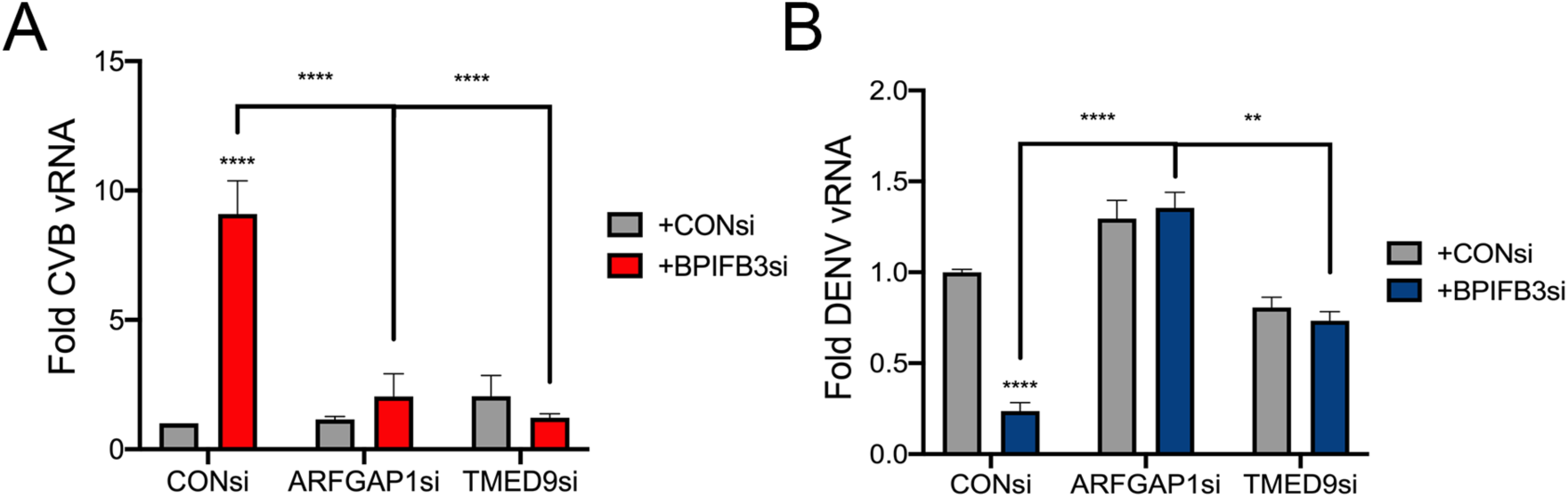
ARFGAP1 and TMED9 reverse the effects of BPIFB3 depletion on viral infection. HBMEC transfected with CONsi or BPIFB3si alone or in combination of ARFGAP1 and TMED9 were infected with CVB (**A**) or DENV (**B**) at a MOI of 1 and analyzed for level of infection by qPCR. Data were analyzed using a two-way ANOVA, ** P < 0.01, **** P <0.0001.

## Discussion

Here, we define the cellular factors that regulate a noncanonical form of autophagy that occurs in the absence of BPIFB3 expression and which is important for RNA virus replication. The characterization of autophagy pathways in mammalian cells is made more complex than the study of parallel pathways in yeast due to significantly increased variability in the signals that control autophagosome biogenesis, the diversity of membranes used for autophagosome formation, and the increased expansion of the ATG8 protein family (Bento et al., 2016). This complexity is further complicated by the collection of autophagy pathways in mammalian cells that serve unique functions, such as organelle specific pathways. Traditionally, macroautophagy functions in response to nutrient stress, specifically amino acid starvation. However, new evidence suggests that changes in lipid content and signals independent from the PI3K complex can play an important role in driving different forms of autophagy (Mishra et al., 2018; Vicinanza et al., 2015; Wang et al., 2015). We have previously identified the ER localized protein BPIFB3 as a negative regulator of a non-canonical autophagy pathway (Delorme-Axford et al., 2014; Evans et al., 2020). While our prior work characterized the induction of autophagy observed during BPIFB3 depletion and eluded to a non-canonical pathway, here we focused on defining the factors required for the induction of BPIFB3-regulated autophagy. The work presented here, demonstrates that many of the regulatory components involved in controlling canonical autophagy are not essential to the form of autophagy induced by decreased BPIFB3 expression. Instead, we show that two host cell factors, ARFGAP1 and TMED9, are required for BPIFB3 induced autophagy, and loss of expression is sufficient to negate the induction of autophagy. We further conclude that ARFGAP1 and TMED9 are required for the effects of BPIFB3 depletion on both CVB and DENV replication. These findings indicate that the enhancement observed in CVB replication is due to the increase in autophagosomes and endosomes within the cytoplasm that occurs during BPIFB3 silencing, which may directly enhance viral replication due to more available membranes for viral RO formation. In contrast, the inhibition of DENV replication that occurs during BPIFB3 silencing, is a result of a disruption in ER integrity and enhanced ER turnover (Evans et al., 2020). Given that these effects are linked to changes in two distinct organelle compartments, our findings support that both ARFGAP1 and TMED9 play essential roles in BPIFB3 function.

BPIFB3 belongs to the BPIFB family of proteins named for their homology to BPI, a secreted antimicrobial protein that functions by binding bacterial lipopolysaccharide (Bingle and Craven, 2002; Bingle et al., 2004; Wong and Levine, 2017). Despite the high degree of predicted structural homology, BPIFB3 and other ER localized members of the family, BPIFB2 and BPIFB6, are not secreted and remain associated with the ER (Delorme-Axford et al., 2014; Morosky et al., 2016). Furthermore, BPIFB proteins are predicted to have a high degree of structural homology to lipid transfer proteins, such as cholesterylester transfer protein (CETP). We previously confirmed that BPIFB3 binds lipids, but have not found direct evidence that it is involved in lipid transfer between membranes like other lipid transfer proteins (Morosky et al., 2016). Despite this, our prior studies have determined that both BPIFB3 and BPIFB6 play important roles in vesicle trafficking, with BPIFB3 impacting autophagy and BPIFB6 disrupting secretory pathway trafficking (Delorme-Axford et al., 2014; Morosky et al., 2016). Recent evidence has demonstrated that the glycolipid transfer protein (GLTP), ceramide-1-phosphate transfer protein (CPTP), plays an essential role in regulating the balance of autophagy induction and apoptosis via direct regulation of Golgi lipid content (Mishra et al., 2018). This mechanism, in agreement with other studies, more broadly implies that the regulation of membrane lipid content alone can directly impact the induction of autophagy or other membrane trafficking events (Chiapparino et al., 2016; Wang et al., 2015). This insight might offer an explanation for how BPIFB3 regulates autophagy independent of each canonical autophagy protein we have tested, and further define how membrane lipid content can directly impact vesicle trafficking events. Beyond the impact of cellular trafficking events, it has been clearly demonstrated that distinct RNA virus families require specific lipid content at sites of RO formation and that virus replication often requires a remodeling of cellular lipids (Albulescu et al., 2015; Chotiwan et al., 2018; Cotton et al., 2017; Strating and van Kuppeveld, 2017; Zhanga et al., 2016). Taken together with the role of lipid transfer proteins in autophagy induction, this suggests a mechanism by which lipid trafficking pathways can directly promote or restrict viral replication.

Given that ARFGAP1 and TMED9 are required for BPIFB3 regulated autophagy and their deletion is sufficient to disrupt this pathway, it is necessary to consider what role these proteins have in the regulation of non-canonical autophagy. ARFGAP1 and TMED9 are both important secretory pathway proteins and have been implicated in the regulation of retrograde COPI vesicle formation (Beck et al., 2009; Strating and Martens, 2009). ARFGAP1 functions as a key GTPase for the well-known secretory pathway GTP binding protein, ARF1. ARFGAP1 has been demonstrated to have important activity in both regulating COPI vesicle formation as well as vesicle coat disassembly (Beck et al., 2009). However, outside of COPI vesicle trafficking, ARFGAP1 has been demonstrated to play a key role in regulating clathrin adapter protein-2 (clathrin-AP-2) endocytosis where it is suggested to function in a similar mechanism to its role in COPI trafficking (Bai et al., 2011), and regulate membrane curvature through its lipid packing sensor (Ambroggio et al., 2010; Spang et al., 2010). In contrast to ARFGAP1’s clearly defined role in vesicle trafficking, TMED9 belongs to the p24 family of proteins which have been broadly implicated in secretory pathway trafficking and COPI vesicle formation, however these processes have not been directly linked to TMED9 (Pastor-Cantizano et al., 2016). Beyond their role in COPI trafficking, there is evidence that both ARFGAP1 and TMED9 effect ER exit sites morphology (Beck et al., 2009; Lavoie et al., 1999), which may be directly tied to their association with BPIFB3 autophagy induction. Many questions remain unanswered, including whether the implicated function of ARFGAP1 and TMED9 within COPI trafficking is related to the role they serve here during BPIFB3si induced autophagy. Our prior studies have shown that loss of BPIFB3 expression has no impact on the localization or morphology of COPI vesicles (Morosky et al., 2016), and the only apparent change we observe are an enhancement of autophagy and disruption of ER integrity (Delorme-Axford et al., 2014; Evans et al., 2020). The data presented here suggest a role for TMED9 in regulating ER exit site morphology during BPIFB3 silencing, however this is not the case upon TMED9 depletion alone, or for ARFGAP1 silencing.

Considering the predicted role of BPIFB3 as a lipid transfer protein, perhaps BPIFB3 functions in lipid trafficking between the ER and Golgi, with ARFGAP1 and TMED9 serving as key regulators of this pathway. In this scenario, we predict depletion of BPIFB3 disrupts the trafficking of lipids, resulting in enhanced autophagy and increased vesicle trafficking out of the ER and directly impacting the replication of RNA viruses that rely on ER derived membranes. While this hypothesis might provide some insight into the function of BPIFB3 and an explanation as to why canonical autophagy regulators are unable to reverse this phenotype, testing this is not trivial. Expression of BPIFB proteins is extremely low and we have yet to be able to detect endogenous BPIFB3 protein to date. Therefore, we have relied on RNAi mediated silencing and ectopic expression of BPIFB3 to study its function. Our results presented here, provide further insight into the requirements for BPIFB3 mediated autophagy and clearly demonstrate ARFGAP1 and TMED9 interact with BPIFB3 to facilitate this pathway.

## Acknowledgements

We thank Terence Dermody (University of Pittsburgh) for providing the BioID2 plasmid and Gerry Hammond (University of Pittsburgh) for providing the VAPB-GFP plasmid. This project was supported by NIH R01-AI081759 (C.B.C.), NIH T32-AI049820 (A.S.E.), a Burroughs Wellcome Investigators in the Pathogenesis of Infectious Disease Award (C.B.C), and the Children’s Hospital of Pittsburgh of the UPMC Health System (C.B.C.).

## Materials and Methods

### Cells and viruses

Human brain microvascular endothelial cells (HBMEC) were maintained in RPMI 1640 supplemented with 10% fetal bovine serum (FBS), 10% NuSerum, 1x non-essential amino acids, 1x minimum essential medium vitamins, 1% sodium pyruvate, and 1% antibiotic. Human bone osteosarcoma U2OS (ATCC HTB-96) and 293T cells were grown in DMEM with 10% FBS and 1% antibiotic. DENV2 16881 was propagated in C6/36 cells (Medina et al., 2012). Titers were determined by fluorescent focus assay as previously described, using recombinant anti-double-stranded RNA monoclonal antibody (provided by Abraham Brass, University of Massachusetts) (Payne et al., 2006). Propagation and titration of CVB3 (RD) has been described previously (Coyne and Bergelson, 2006). All experiments measuring infection levels were performed using a multiplicity of infection (MOI) of 1 for 16 hours (CVB) or 48 hours (DENV) unless stated otherwise, and infection was quantified by RT-qPCR.

### BioID2 assay

HBMEC were seeded in 2 10 cm dishes and transiently transfected with BPIFB3-BioID2 or MCS-BioID2. 24 hours post transfection, cells were rinsed with PBS and media was replaced containing 50 μM D-Biotin and incubated at 37 C for 24 hours. Media was removed and washed twice with PBS. Cells were lysed using 1X RIPA lysis buffer at pH 7.4 and incubated with monomeric Avidin resin overnight (∼16 hours). Purification of biotinylated proteins was performed according to the previously described protocol (Kim et al., 2016). Purified protein extracts were submitted to MSBioworks for further purification and mass spectrometry analysis.

### siRNAs, plasmids and transfections

Characterization of siRNA targeting BPIFB3 was described previously and CONsi and BPIFB3si werw purchased from Sigma (Delorme-Axford et al., 2014). ON-TARGETplus SMARTpool siRNAs targeting ARFGAP1 and TMED9 were purchased from Dharmacon. Sequences for pooled siRNAs are as follows: ARFGAP1 (GAGAGGAGGAGCUCGGACA, CAGGAUGAGAACAACGUUU, GCCACAGCCUGAACGAGAA, CGUCCAUGGUGCACCGAGU), TMED9 (GGACGCAGCUGUAUGACAA, CGGGCUGGGUAGAGUGAUG, AGUGCUUUAUUGAGGAGAU, ACAUCGGAGAGACGGAGAA). Efficiency of knockdown was determined by RT-qPCR for each siRNA target. siRNAs were reverse transfected at 25 nM in to HBMEC using Dharmafect 1, and cells were either infected or RNA was collected 48 hrs post transfection.

Development of GFP tagged ARFGAP1 and TMED9 was accomplished according to the manufacturers protocol using the CT-GFP Fusion TOPO™ Expression Kit purchased from ThermoFisher Scientific. VAPB-eGFP was a gift from Dr. Gerry Hammond (University of Pittsburgh). Plasmids were transfected into U2OS and 293T cells using X-tremeGENE™ 9 DNA Transfection Reagent (Sigma) or HBMEC cells using Lipofectamine 3000 (ThermoFisher) according to the manufacturers protocols and fixed for fluorescence microscopy or lysed for immunoprecipitation 48 hrs post transfection.

### RNA extraction, cDNA synthesis, and RT-qPCR

RNA was isolated using the GenElute Total RNA MiniPrep kit from Sigma according to the kit protocol. RNA was reverse transcribed using the iScript cDNA Synthesis kit (Bio-Rad) with 1μg of RNA per sample. RT-qPCR was performed using IQ SYBR green SuperMix (Bio-Rad) in a Bio-Rad CFX96 Touch real-time PCR detection system. A modified threshold cycle (ΔCT) method was used to calculate gene expression using human actin for normalization. Primer sequences for actin, DENV, CVB, and BPIFB3 have been described previously (Delorme-Axford et al., 2013; Lennemann and Coyne, 2017). Predesigned KiCqstart qPCR primers for ARFGAP1 (H_ARFGAP1_1) and TMED9 (H_TMED9_1) were purchased from Sigma.

### Antibodies

Mouse monoclonal anti-V5 epitope tag was purchased from Invitrogen (R960-25). Rabbit anti-LC3B (ab48394) and mouse anti-p62 (ab56416) were purchased from Abcam. Mouse monoclonal antibody against LAMP1 (H4A3) and rabbit monoclonal antibody against GAPDH (FL-335) were purchased from Santa Cruz Biotechnology. Mouse monoclonal Sec31A (612350) antibody was purchased from BD Biosciences and Alexa Fluor conjugated secondary antibodies were purchased from Invitrogen.

### Immunofluorescence and electron microscopy

Immunofluorescence microscopy was performed on cells grown in 8-well chamber slides (Millipore Sigma), fixed in 4% paraformaldehyde, and permeabilized with 0.1% Triton. Primary antibodies were incubated in PBS with cells for 1 hr, followed by staining with Alexa Fluor conjugated secondary antibodies for 30 min. Slides were mounted with coverslips using VectaShield containing 40-6-diamino-2-phenylindole (DAPI). Imaging was performed on an Olympus IX83 inverted microscope or a Zeiss LSM 710 confocal microscope. All image quantification was performed using ImageJ/FIJI. Quantification of fluorescent puncta was performed manually. Preparation of samples for TEM were done as previously described, by the Center for Biologic Imaging (University of Pittsburgh) (Delorme-Axford et al., 2013). Imaging was performed on a JEM 1400Plus transmission electron microscope. Quantification of TEM images was performed manually.

### Statistical analyses

All analyses were performed using GraphPad Prism. Experiments were performed at least three times. Student’s t test, 2way analysis of variance (ANOVA), or one-way ANOVA were used where indicated. Analysis of fluorescent microscopy data was done using a non-parametric Kruskal-Wallis test. Data are presented as mean ± standard deviation, with specific p-values detailed in the figure legends.

## Supplemental Figures

**Figure S1:**
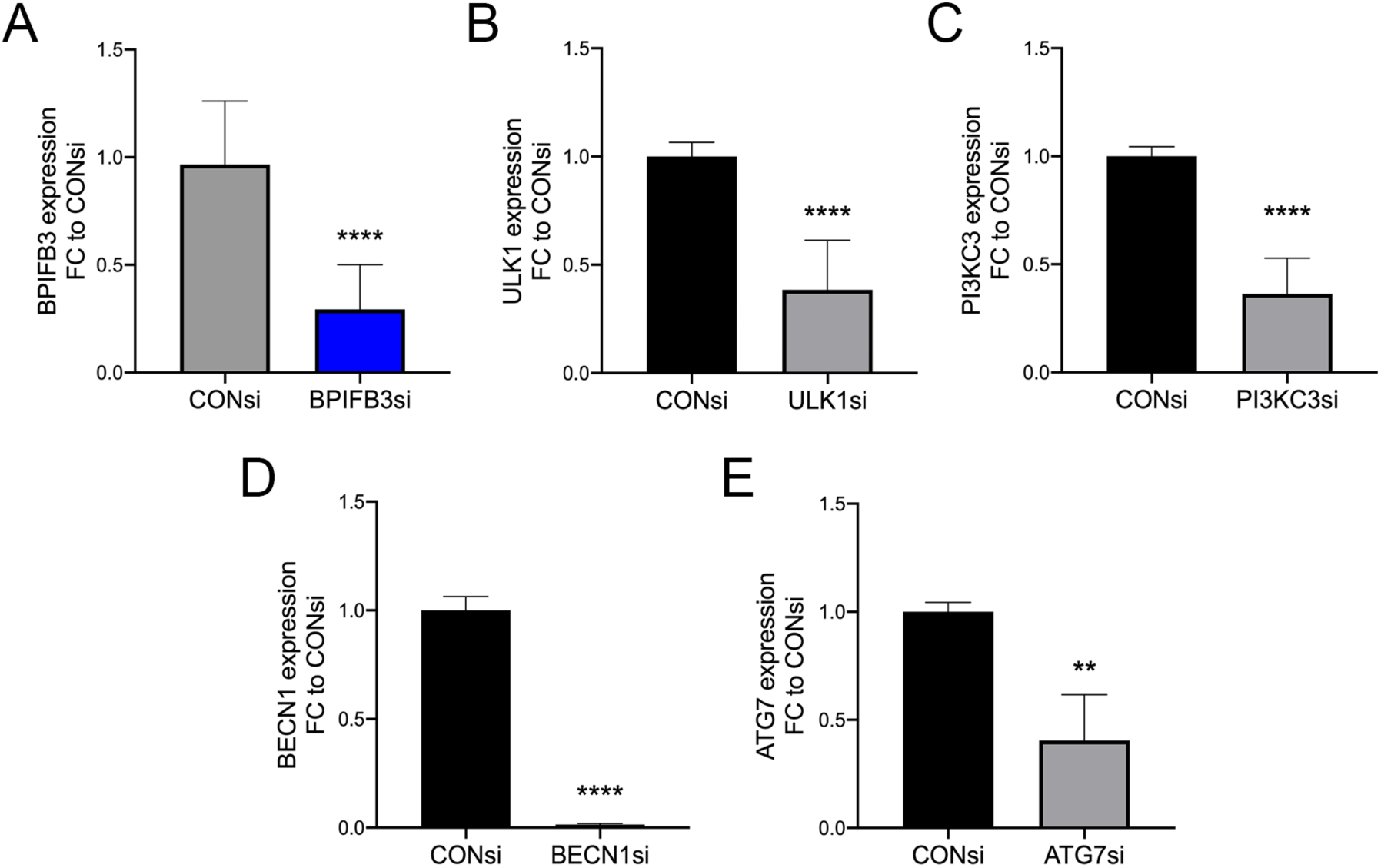
siRNA knockdown efficiency of autophagy regulatory components. The knockdown efficiency for each siRNA targeting BPIFB3 (A), ULK1 (B), PI3KC3 (C), BECN1 (D), and ATG7 (E) was determined by qPCR for target mRNA and expressed as fold change to control siRNA transfected cells (CONsi). Data were analyzed using an unpaired t test, ** P < 0.01, **** P < 0.0001.

**Figure S2:**
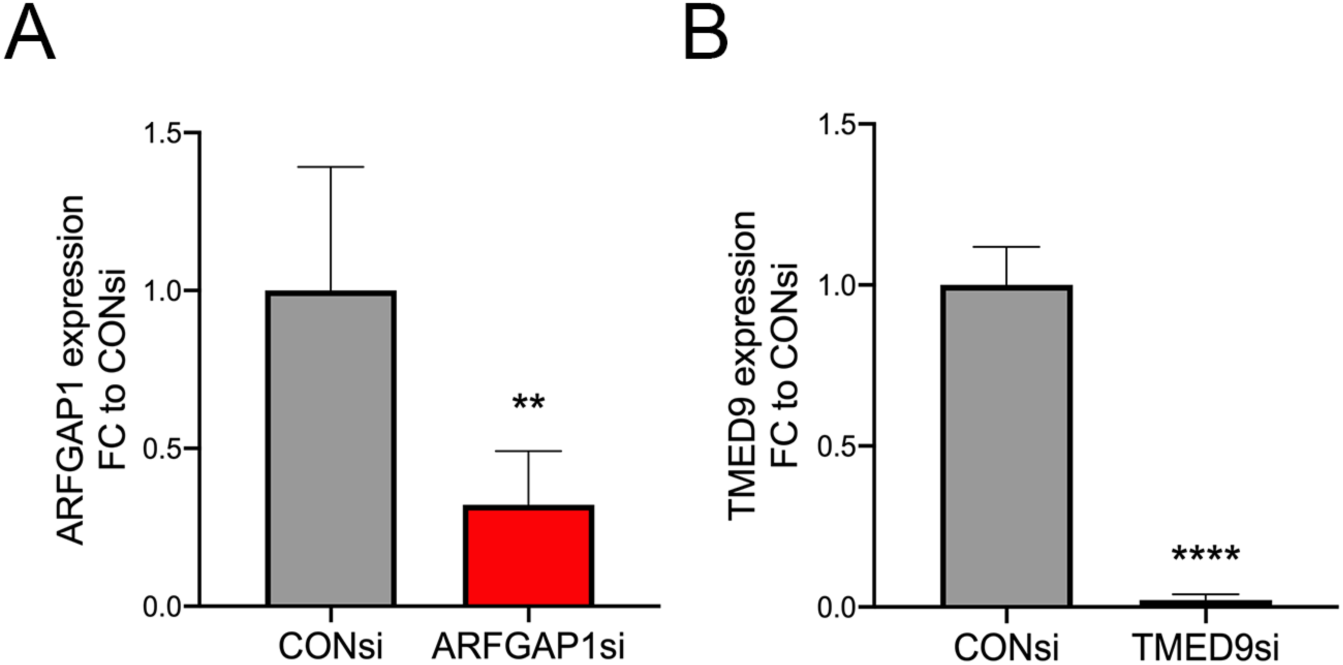
siRNA knockdown efficiency and ARFGAP1 and TMED9. The knockdown efficiency for siRNAs targeting (A) ARFGAP1, and (B) TMED9 was determined by qPCR for target mRNA and expressed as fold change to control siRNA transfected cells (CONsi). Data were analyzed using an unpaired t test, ** P < 0.01, **** P < 0.0001.

